# Adaptation and Selection Shape Clonal Evolution During Residual Disease and Recurrence

**DOI:** 10.1101/2020.04.22.055780

**Authors:** Andrea Walens, Jiaxing Lin, Jeffrey S. Damrauer, Ryan Lupo, Rachel Newcomb, Douglas B. Fox, Nathaniel W. Mabe, Jeremy Gresham, Tristan De Buysscher, Hemant Kelkar, Piotr A. Mieczkowski, Kouros Owzar, James V. Alvarez

## Abstract

The survival of residual tumor cells following therapy and their eventual recurrence constitutes one of the biggest obstacles to obtaining cures in breast cancer, but it remains unclear how the clonal composition of tumors changes during tumor relapse. We used cellular barcoding to directly monitor clonal dynamics during tumor recurrence in a genetically engineered mouse model. We found that the clonal diversity of tumors progressively decreased during tumor regression, residual disease, and recurrence. Only a fraction of subclones survived oncogene withdrawal and persisted in residual tumors. The minimal residual disease phase itself was accompanied by a continued attrition of clones, suggesting an ongoing process of selection during dormancy. The reactivation of dormant residual cells into recurrent tumors followed several distinct evolutionary routes. Approximately half of the recurrent tumors exhibited a striking clonal dominance in which one or two subclones comprised the vast majority of the tumor. The majority of these clonal recurrent tumors exhibited evidence of *de novo* acquisition of Met amplification, and were sensitive to small-molecule Met inhibitors. A second group of recurrent tumors exhibited marked polyclonality, with thousands of subclones and a clonal architecture very similar to primary tumors. These polyclonal recurrent tumors were not sensitive to Met inhibitors, but were instead dependent upon an autocrine IL-6 – Stat3 pathway. These results suggest that the survival and reactivation of dormant tumors can proceed via multiple independent routes, producing recurrent tumors with distinct clonal composition, genetic alterations, and drug sensitivities.

## Introduction

Tumor recurrence after initial treatment is a frequent cause of death in many cancers, including breast cancer, and recurrent tumors are often resistant to therapies to which the corresponding primary tumors were sensitive (Acharyya et al., 2012; Gonzalez-Angulo et al., 2007; Jones, 2008; Saphner et al., 1996). Recurrent tumors are thought to arise from a population of cells that survives initial treatment; these cells, often referred to as minimal residual disease, can persist in a clinically undetectable state for years or even decades before resuming growth to give rise to recurrent tumors (Goss and Chambers, 2010; Klein, 2011; Sosa et al., 2014). In spite of the clinical importance of residual disease and recurrence, little is known about the pathways that regulate the longterm survival of residual cells or that induce the reactivation of these cells to yield recurrent tumors (Bivona and Doebele, 2016). Identifying such pathways may suggest opportunities for directly killing residual cells or preventing their reactivation, thereby forestalling the development of recurrent tumors (Ghajar, 2015).

Primary breast tumors are heterogeneous, harboring different subclones of genetically or epigenetically distinct cells (Marusyk et al., 2012). While tumor progression is thought to be driven by the progressive outgrowth of aggressive subclones, little is known about how the clonal composition of tumors changes during residual disease and recurrence. Understanding the clonal dynamics of dormancy and recurrence is essential for developing strategies to prevent or treat recurrence. However, studying these processes in patients is challenging, given the difficulty in identifying residual disease and obtaining recurrent tumors.

To overcome these obstacles, we and others have used an inducible genetically engineered mouse model that exhibits key features of breast cancer progression, including the survival of cancer cells following therapy and their eventual spontaneous recurrence (Abravanel et al., 2015; Alvarez et al., 2013; Feng et al., 2014; Mabe et al., 2018; Moody et al., 2005; Moody et al., 2002). In this model, doxycycline (dox) administration to MMTV-rtTA;TetO-neu (MTB;TAN) mice induces expression of the Her2/neu oncogene, leading to the formation of invasive mammary tumors. Subsequent withdrawal of dox induces Her2 downregulation and complete tumor regression, perhaps mimicking anti-Her2 targeted therapy. However, a small population of residual cells survives Her2 downregulation and persists in a dormant, non-proliferative state in the mammary gland (Feng et al., 2014). These residual cells eventually re-initiate proliferation, independent of Her2 expression, to form a recurrent tumor. Here we combined this conditional mouse model with lentiviral-mediated cellular barcoding to study the clonal dynamics of tumor regression, residual disease and relapse.

## Results

### Cellular barcoding to track clonal dynamics during tumor regression, residual disease, and recurrence

We used a cellular barcoding strategy to directly monitor changes in the clonal composition of tumors during tumor regression, residual disease, and recurrence. In this approach, random, inert DNA barcodes are introduced into a population of cells. Each barcode serves as a molecular tag that marks all of the progeny of a particular cell; all cells marked by a particular barcode are defined here as a clone, reflecting the fact that they arose from a single cell. The clonal composition of a population can be determined by measuring the number of clones (number of unique barcodes detected) and the abundance of each clone (proportion of total reads contributed by each barcode) using next-generation sequencing. In this manner, it is possible to track the clonal dynamics of a cell population under different conditions. This approach has been used to study clonal dynamics in response to targeted therapies in vitro (Bhang et al., 2015; Hata et al., 2016) and during the growth of tumors in vivo (Nguyen et al., 2014; Nguyen et al., 2015).

To implement this barcoding strategy we digested a primary Her2-driven tumor (donor tumor #1) from an MTB;TAN mouse and cultured tumor cells ex vivo in the presence of dox to maintain Her2 expression. Approximately 200,000 tumor cells were then infected with a lentiviral barcode library at a multiplicity of infection of 0.1 to ensure that each cell received a single DNA barcode. Following selection, the population was expanded for 12 population doublings, yielding a population of 100 million cells containing approximately 20,000 unique barcodes with each barcode represented in an average of 5,000 cells (Figure 1A). This barcoded cell population was injected (1 million cells per injection, 50 cells per barcode) into the mammary glands of a large cohort of recipient mice on dox (Figure 1B). Primary tumors formed approximately 3 weeks following injection (Figure 1C). Once primary tumors reached 5 mm in diameter, one cohort of mice was sacrificed with primary tumors (n=6 tumors), and the remaining mice were removed from dox to induce Her2 downregulation, leading to rapid tumor regression (Figure 1C). Additional cohorts of mice were sacrificed with residual tumors at 4 and 8 weeks following dox withdrawal (n=6 tumors per cohort), prior to the time point at which most mice develop recurrent tumors (Figure 1C and D). A final cohort of mice was allowed to develop recurrent tumors (n=12 tumors); recurrent tumors in these mice arose with a median time of 75 days (Figure 1D). Importantly, this timing is similar to mice injected with primary tumor cells lacking DNA barcodes (data not shown), suggesting that introduction of this barcode library does not affect the timing of recurrent tumor formation.

**Figure 1.**
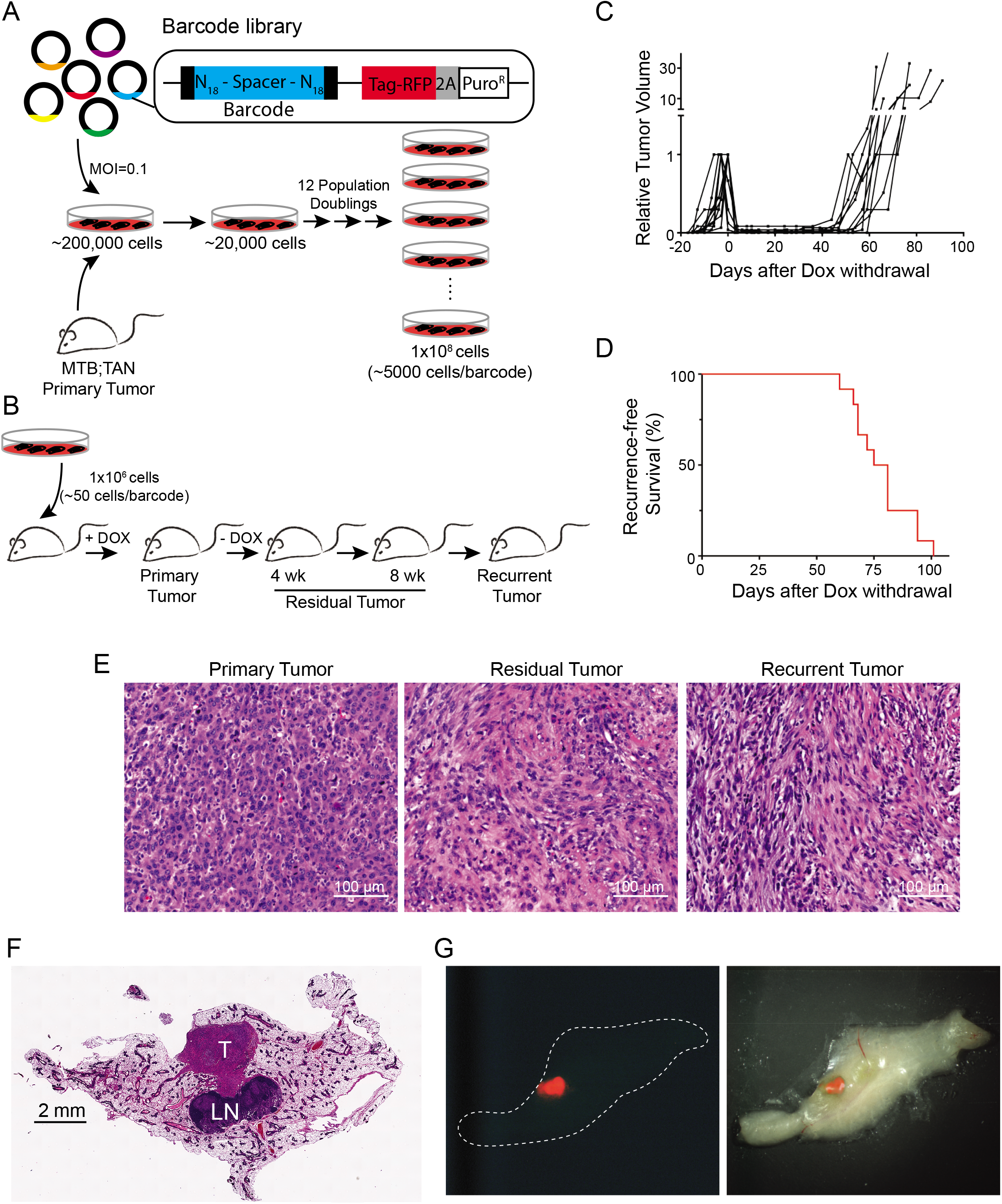
Cellular Barcoding to Track Clonal Dynamics During Tumor Regression, Residual Disease, and Recurrence. (A) Schematic of cellular barcoding strategy. Primary Her2-driven mouse mammary tumors were digested, cultured with Dox, and 200,000 cells were infected at low MOI (0.1) with a lentiviral barcode library comprising ~60 million unique barcodes. Following selection for transduced cells, the population was expanded to yield a population of 100 million cells comprising ~20,000 unique barcodes (~5000 cells/barcode). (B) One million barcoded tumor cells were injected bilaterally into the inguinal mammary glands of nu/nu mice on Dox. Once orthotopic primary tumors reached 5 mm in diameter, one cohort of mice was sacrificed with primary tumors (n=6 tumors). Dox was removed from the remaining mice, and cohorts were sacrificed with residual tumors after 4 weeks (n=6 tumors) and 8 weeks (n=6 tumors). A final cohort of mice was monitored until recurrent tumors formed (n=12 tumors). (C) Tumor growth curves showing primary tumor growth, regression, residual disease, and recurrence for representative orthotopic Her2-driven tumors. (D) Kaplan-Meier recurrence-free survival curves for barcoded orthotopic tumors. (E) H&E-stained sections of a representative primary, residual, and recurrent tumor. Scale bar = 100 μm. (F) H&E-stained section of a mammary gland whole-mount showing the size and location of a representative residual tumor. T = tumor, LN = lymph node. Scale bar = 2 mm. (G) Fluorescent (left) and merged fluorescent-brightfield (right) image of a representative fluorescently labeled residual tumor (red) within the mammary gland.

Primary and recurrent tumors arising in this orthotopic setting resemble histologically tumors from the autochthonous model. Primary tumors exhibited an epithelial morphology, with nests of epithelial cells surrounded by stromal cells (Figure 1E, left). Recurrent tumors had a mesenchymal morphology, consistent with the previous finding that tumor recurrence in these models is associated with epithelial-to-mesenchymal transition (Moody et al., 2005) (Figure 1E, right). Interestingly, residual tumors comprised cells with both epithelial and mesenchymal characteristics, and also had a notable stromal component (Figure 1E, middle).

To measure the barcode composition of these tumors, we next isolated genomic DNA from tumors at each time point. To ensure accurate representation of barcodes, we isolated genomic DNA from the entire tissue sample for both primary and recurrent tumors. Although residual tumors were readily apparent on an H&E stained section (Figure 1F), they were too small to identify grossly. Therefore, to facilitate isolation of barcodes from these residual tumors, tumors were microdissected using a fluorescence microscope prior to DNA isolation (Figure 1G). In this manner, we were able to isolate barcoded genomic DNA from primary, residual, and recurrent tumors for subsequent sequencing.

### Primary tumors are driven by the expansion of a subset of clones

We used next-generation sequencing to measure the number of barcodes and their relative abundance in each tumor. The starting cell population contained approximately 17,000 unique barcodes (Figure 2A and B), consistent with our target of 20,000 barcodes. The mean barcode frequency was 0.00588%, and the frequency of the most abundant barcode was 0.707% (range: 0.0000736% – 0.707%; Figure 2A). This suggests that the growth of these cells in vitro did not impose a strong selection for individual clones. In contrast, an average of 4,544 barcodes were detected in primary tumors (range: 3,063 to 5,648), and there was evidence for a selection for individual barcodes (Figure 2A and B). To quantitatively assess the clonal complexity of each tumor we used the Shannon Diversity index (Magurran, 2005), a measure of the diversity of a population that incorporates both the number of clones and their representation. Tumors with many clones that are evenly distributed have a high Shannon Index, while tumors with few or dominant clones have a low Shannon Index. This analysis revealed that primary tumors had a reduction in barcode complexity as compared to the starting population (Figure 2C). Consistent with this, barcodes were more unevenly distributed in primary tumors: 50% of reads were contributed by an average of 56 barcodes in primary tumors, as compared to 700 barcodes in the starting population (Figure 2D). The most abundant barcode in primary tumors represented an average of 8.4% of the population (range: 4.5-12.8%), as compared to 0.707% in the starting population (Figure 2E-G and Supplemental Figure 1). Taken together, these results suggest that primary tumor formation is accompanied by a reduction in clonal complexity and the expansion of a subset of clones.

**Figure 2.**
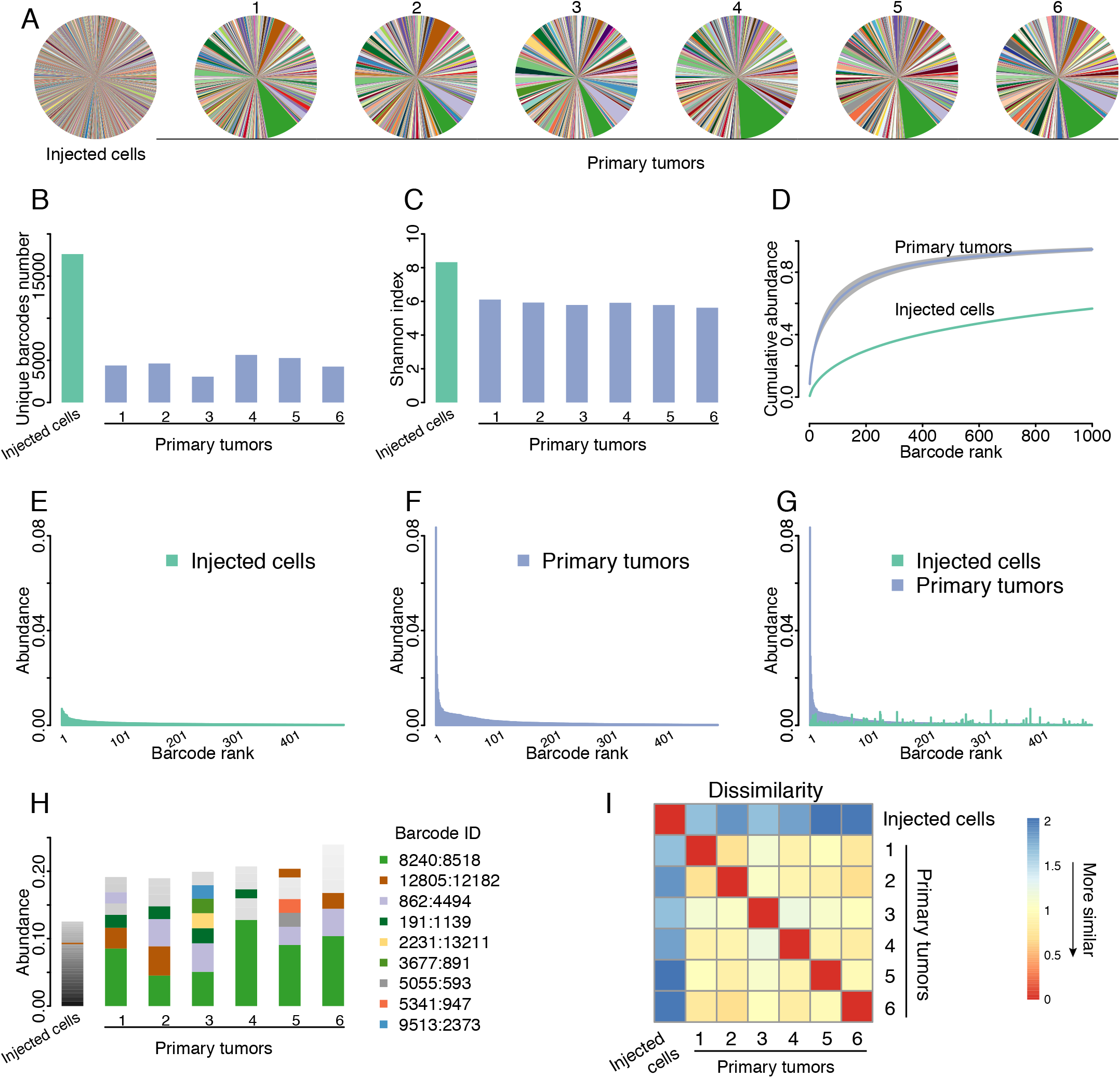
Independent Primary Tumors Have Similar Clonal Composition. (A) Pie charts showing the relative frequency of barcodes in the starting cell population and 6 independent primary tumors. Note that individual barcodes are represented by the same color in each pie chart. (B) Number of unique barcodes detected in the starting cell population and 6 independent primary tumors. (C) Shannon diversity index showing barcode complexity of the starting cell population and 6 independent primary tumors. (D) Cumulative abundance plots for the starting cell population and the average of 6 primary tumors, demonstrating that barcode complexity decreases during primary tumor growth. (E) Barcode abundance in the starting cell population. Barcodes are ranked on the x-axis from most to least abundant. (F) Barcode abundance in a representative primary tumor. Barcodes are ranked on the x-axis from most to least abundant. (G) Comparison of barcode distribution in the starting cell population and a representative tumor, showing that the most abundant barcodes in tumors were not abundant in the starting population. Barcodes are ranked on the x-axis based on their abundance in the primary tumor. (H) A subset of barcodes is reproducibly enriched in independent primary tumors. The most abundant barcodes in each sample are shown, with each color denoting a unique barcode. Note that barcode colors match pie charts in (A). (I) Correlation matrix showing the similarity in barcode abundance between samples. Primary tumors had similar barcode distributions to one another, but were distinct from the starting population. The Jensen-Shannon divergence was used to measure dissimilarity among tumors.

### Independent primary tumors have similar clonal composition

The observation that the formation of primary tumors is associated with a reduction in clonal complexity suggests that tumor growth in vivo imposes different or more stringent selective pressures than growth in vitro. We considered two possibilities: abundant clones in primary tumors may have also been abundant in the starting population and simply expanded further during tumor growth in vivo. Alternatively, tumor growth may have selected for a distinct subset of clones that were not abundant in the starting population. To distinguish among these possibilities, we compared abundant clones between tumors and the starting population. The barcodes that were most abundant in primary tumors were not abundant in the injected cell population (Figure 2G), suggesting that tumor growth in vivo selects for distinct clones. We next explored whether the same clones were most abundant in independent primary tumors, which would suggest that there were pre-existing clones in the starting population with an inherent capacity to grow in vivo. Examination of the barcodes composing the top 15% of reads revealed that the same set of barcodes were most abundant in independent primary tumors, but not in the staring population of cells (Figure 2H). For instance, barcode 8240:8518 was the most abundant in all primary tumors, and represented an average of 8.4% of total reads in primary tumors (range: 4.5-12.8%). In contrast, this barcode composed only 0.027% of reads in the starting cell population, indicating that the clone marked by this barcode expanded 300-fold during primary tumor growth. To quantitatively assess the similarity in clonal composition of tumors we calculated the Jensen-Shannon divergence (Fuglede and Topsoe, 2004) between barcode abundances in independent primary tumors. All primary tumors were highly similar to one another but dissimilar from the starting population of injected cells (Figure 2I). Taken together, these results indicate that primary tumor growth selects for a subset of pre-existing clones with increased tumorigenic potential, and highlights the utility of cellular barcoding to identify such clones.

### Reduction in clonal complexity during tumor regression and residual disease

We next asked how the clonal composition of tumors changed following tumor regression and during residual disease. As described above, Her2 downregulation in primary tumors leads to near complete tumor regression, but a small population of tumor cells survives and persists in a non-proliferative state (see Figure 1F and G). Tumors were harvested at 4 weeks (early residual tumors) and 8 weeks (late residual tumors) following dox withdrawal. Comparing primary tumors to early residual tumors provides insight into changes in clonal composition during tumor regression, while comparisons between early and late residual tumors can reveal changes that occur during residual disease.

Sequencing of barcodes in tumors 4 weeks after dox withdrawal revealed that tumor regression was accompanied by a decrease in the clonal complexity of tumors and the further enrichment of a subset of clones (Figure 3A). Early residual tumors had fewer barcodes (Figure 3C; P-value = 2.6× 10^-3^) and a reduction in barcode complexity (Figure 3D; P-value = 4.8× 10^-4^) as compared to primary tumors. Barcodes in these tumors were more unevenly distributed than primary tumors, with 50% of total reads coming from only ~16 barcodes (Figure 3E; P-value = 3.3 × 10^-4^). Barcode 8240:8518 – which was the most abundant in primary tumors – was further enriched following tumor regression, and composed on average of 17.4% of the population (range: 4.2-34.1%; Figure 3A and Supplemental Figure 2). Taken together, these results indicate that tumor regression following Her2 downregulation is accompanied by a reduction in clonal complexity.

**Figure 3.**
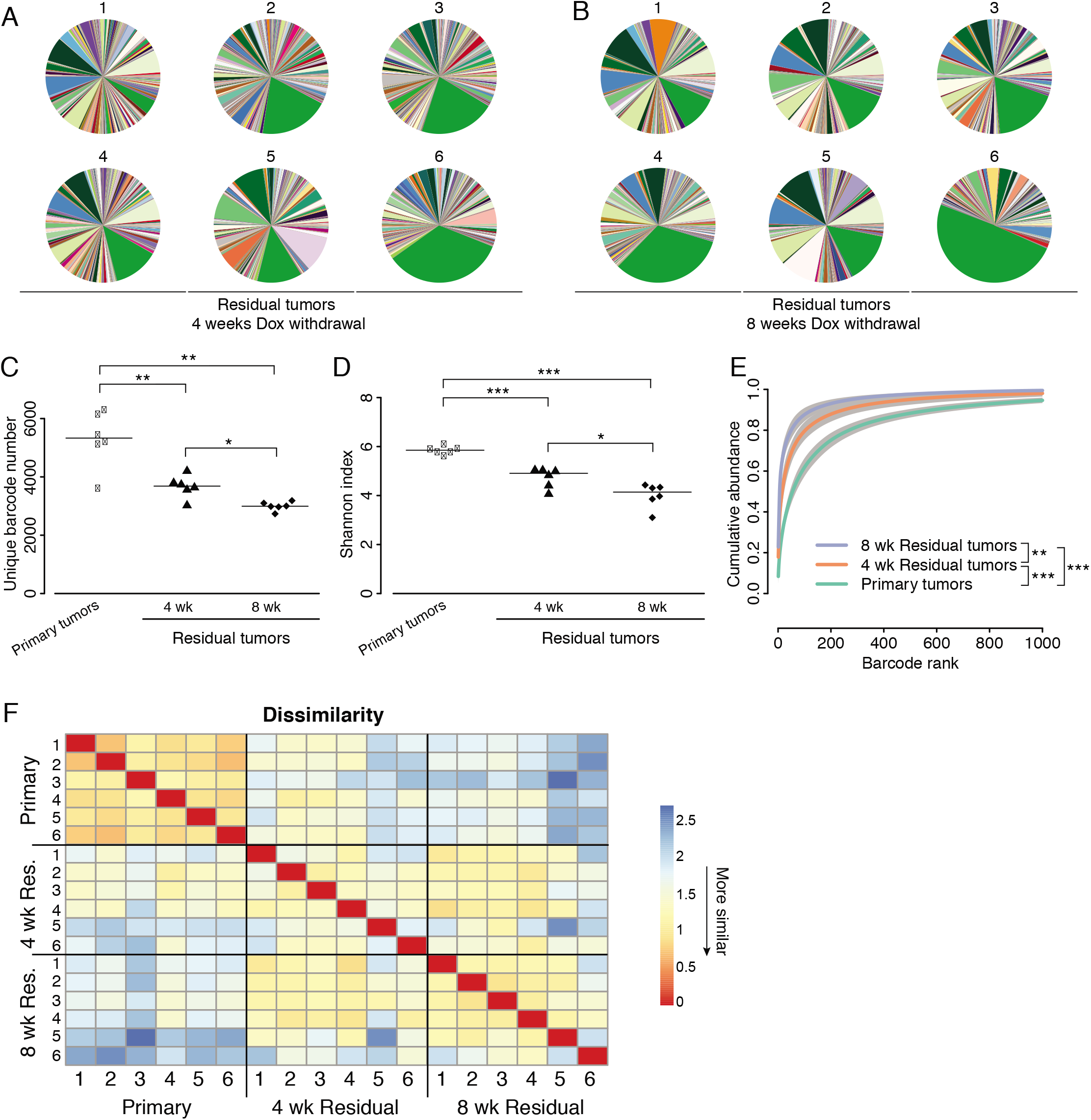
Clonal Complexity Decreases During Tumor Regression and Residual Disease. (A) Pie charts showing the relative frequency of barcodes in early residual tumors (4 weeks following Dox withdrawal). (B) Pie charts showing the relative frequency of barcodes in late residual tumors (8 weeks following Dox withdrawal). (C) Number of unique barcodes detected in primary tumors and residual tumors 4 and 8 weeks following Dox withdrawal. * P < 0.05, ** P < 0.01. Significance determined by Welch Two Sample T-test. (D) Shannon Diversity Index showing barcode complexity in primary tumors and residual tumors 4 and 8 weeks following Dox withdrawal. * P < 0.05, *** P < 0.001. Significance determined by Welch Two Sample T-test. (E) Cumulative abundance plots for primary tumors and residual tumors 4 and 8 weeks following Dox withdrawal. Barcode complexity decreased following tumor regression and continued to decrease during the persistence of residual disease. ** P < 0.01, *** P < 0.001. Significance determined by Welch Two Sample T-test between the number of barcodes composing 50% of reads. (F) Correlation matrix showing the similarity in barcode abundance between samples demonstrating that the barcode distribution progressively changes during residual disease. The Jensen-Shannon divergence was used to measure dissimilarity among tumors.

Following tumor regression, residual tumors persist for between one and two months before developing into recurrent tumors. We next determined how the clonal composition of tumors changes during this residual disease period by comparing barcode number and abundance between early and late residual tumors. Surprisingly, we found that there was a continued reduction in clonal complexity during this residual disease stage (Figure 3A, B). Late residual tumors had fewer barcodes (P-value = 2.4×10^-2^), reduced Shannon index (P-value = 2.0×10^-2^), and a more uneven distribution of barcodes (P-value = 8.0×10^-3^) as compared to early residual tumors, with 50% of total reads coming from 5.5 barcodes on average (Figure 3C-E). Barcode 8240:8518 remained the most abundant barcode in all late residual tumors, and represented as high as 49% of total reads (range: 10.4-49.2%; Figure 3A and Supplemental Figure 2). Notably, the decrease in clonal complexity between 4 and 8 weeks post-dox withdrawal was similar in magnitude to the decrease in complexity that accompanied tumor regression (Figure 3D). This suggests that the residual disease state is accompanied by a continued attrition of a subset of clones, perhaps indicative of ongoing selective pressures during residual disease.

To gain insight into how the overall clonal architecture of tumors changed during tumor regression and residual disease, we examined the similarity of barcode abundances between residual tumors and primary tumors based on the Jensen-Shannon divergence. As shown above, primary tumors were highly similar to one another (Figure 2I and 3F). In contrast, the barcode composition of early and late residual tumors was progressively less similar to that of primary tumors (Figure 3F). In addition, residual tumors were more dissimilar from one another than primary tumors were from one another (Figure 3F). Taken together, these results suggest that the clonal composition of tumors progressively changes during tumor regression and residual disease, with residual tumors exhibiting a greater divergence in their clonal composition.

### Distinct clonal architecture in recurrent tumors suggests divergent routes to recurrence

The findings above indicate that tumor regression and residual disease were accompanied by a decrease in the number and complexity of clones. Nonetheless, all residual tumors still retained several thousand clones that could serve as a template for further selection. We reasoned that tumor recurrence could be driven by the reactivation of a small subset of clones, or by a broader, tumor-wide reactivation of many or most of the clones in residual tumors. To distinguish between these possibilities, we examined the clonal composition of recurrent tumors. We found that recurrent tumors exhibited a surprisingly large variation in the number and distribution of barcodes (Figure 4A). Three recurrent tumors (recurrent tumors #1-3) contained thousands of barcodes that were relatively evenly distributed, as evidenced by a high diversity index (Figure 4A-B). At the other end of the spectrum, in three recurrent tumors (recurrent tumors #10-12) almost all reads were contributed from a single barcode (Figure 4A). The grossly uneven distribution of barcodes in these clonal tumors was reflected by a very low diversity index (Figure 4B). Interestingly, in recurrent tumor #10 and #12, the most abundant barcode, 8240:8518, was also the most abundant barcode in primary and residual tumors (Figure 4A and Supplemental Figure 2). The remaining tumors (recurrent tumors #4-9) had an intermediate clonal complexity (Figure 4A-B). Tumors #4, 6, 8, and 9 had between two and eight dominant barcodes, while tumors #5 and 7 had a single abundant barcode together with a large number of minor barcodes.

**Figure 4.**
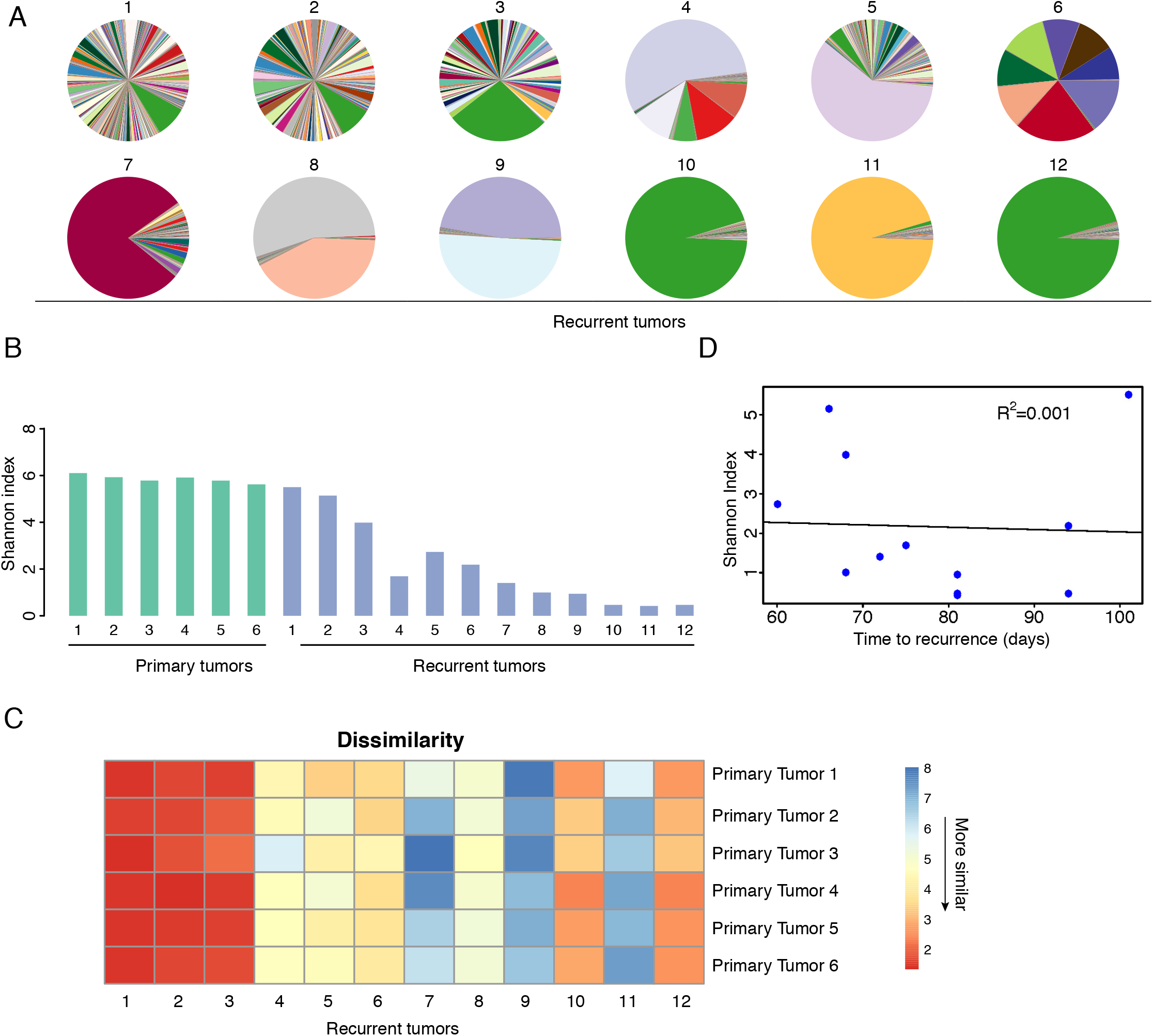
Distinct Clonal Composition in Recurrent Tumors. (A) Pie charts showing the relative frequency of barcodes in recurrent tumors. (B) Shannon Diversity Index showing barcode complexity in primary and recurrent tumors. (C) Correlation matrix showing the similarity in barcode abundance between recurrent tumors and primary tumors. Recurrent tumors fall into three groups, one group (#1-3) whose barcode distribution is highly similar to primary tumors, one group (#10 and 12) with intermediate similarity, and one group (#4-9, 11) whose barcode distribution is very dissimilar. The Jensen-Shannon divergence was used to measure dissimilarity among tumors. (D) Barcode diversity in recurrent tumors is not correlated with time to recurrence.

We next assessed how the clonal architecture of these recurrent tumors compared to primary tumors. The barcode distribution of tumors #1-3 was very similar to primary tumors, suggesting that the clonal composition of these recurrent tumors closely resembled that of primary tumors (Figure 4C). Tumors #10 and 12 had intermediate similarity to primary tumors; this was influenced by the fact that barcode 8240:8518 was the most abundant barcode in these tumors (Figure 4D and Supplemental Figure 2). In contrast, the remaining recurrent tumors were very dissimilar from primary tumors (Figure 4C), indicating that these tumors were dominated by distinct clones from those present in primary tumors.

Finally, we asked whether clonal and polyclonal tumors recurred with different kinetics. We found that there was no correlation between the clonal diversity of recurrent tumors and their time to recurrence (Figure 4D). In sum, these results suggest that recurrence can proceed through at least two distinct routes. One route proceeds through a tumor-wide reactivation of all or most clones in a residual tumor, yielding a recurrent tumor with thousands of clones whose distribution resembles primary tumors. In the second route, only one or a few clones resume growth, giving rise to (oligo)clonal tumors dominated by clones distinct from those found in primary tumors.

### Met amplification drives (oligo)clonal recurrences that are sensitive to Met inhibitors

We next wanted to understand the mechanistic basis for the different clonal architecture found in recurrent tumors. Signaling through the receptor tyrosine kinase c-Met has been shown to be a common escape mechanism for tumors following loss of oncogenic signaling. For instance, Met amplification has been shown to promote resistance to EGFR inhibitors in EGFR mutant non-small-cell lung cancer (Bhang et al., 2015; Engelman et al., 2007; Turke et al., 2010). In a mammary tumor model with conditional PIK3CA expression, Met amplification drove tumor recurrence following PIK3CA downregulation (Liu et al., 2011). Finally, increased signaling through Met was shown to promote recurrence in the same Her2-driven model we are using here (Feng et al., 2014). We therefore tested whether Met amplification occurred in a subset of recurrent tumors, and whether differences in Met amplification status could underlie the different clonal composition of recurrent tumors. We measured Met copy number using a qPCR-based copy-number assay on genomic DNA, and found that 6 of the 12 recurrent tumors had Met amplification (Figure 5A). Interestingly, the most abundant barcode(s) in each Met-amplified tumor was different (Figure 5B), suggesting that these Met-amplified clones are distinct from one another. Consistent with this, mapping of the amplicon boundaries using qPCR on neighboring genes revealed that different recurrent tumors had distinct amplicons (Figure 5C), suggesting that these Met-amplified tumors arose from different Met amplified clones. Surprisingly, the barcodes marking these Met-amplified clones could be detected in all other recurrent tumors – as well as in all primary and residual tumors – but at much lower frequencies (Supplemental Figure 3A-G). This suggests either that Met amplification is not by itself sufficient for recurrence, or that Met amplification occurs de novo in each tumor. In EGFR-mutant lung cancer, populations of cells that have pre-existing resistant clones develop resistance more quickly than populations where resistance develops de novo (Hata et al., 2016). We therefore compared the recurrence time between Met-amplified and non-amplified tumors. Surprisingly, we could detect no difference in recurrence-free survival between these tumors (Figure 5D; P-value = 0.61).

**Figure 5.**
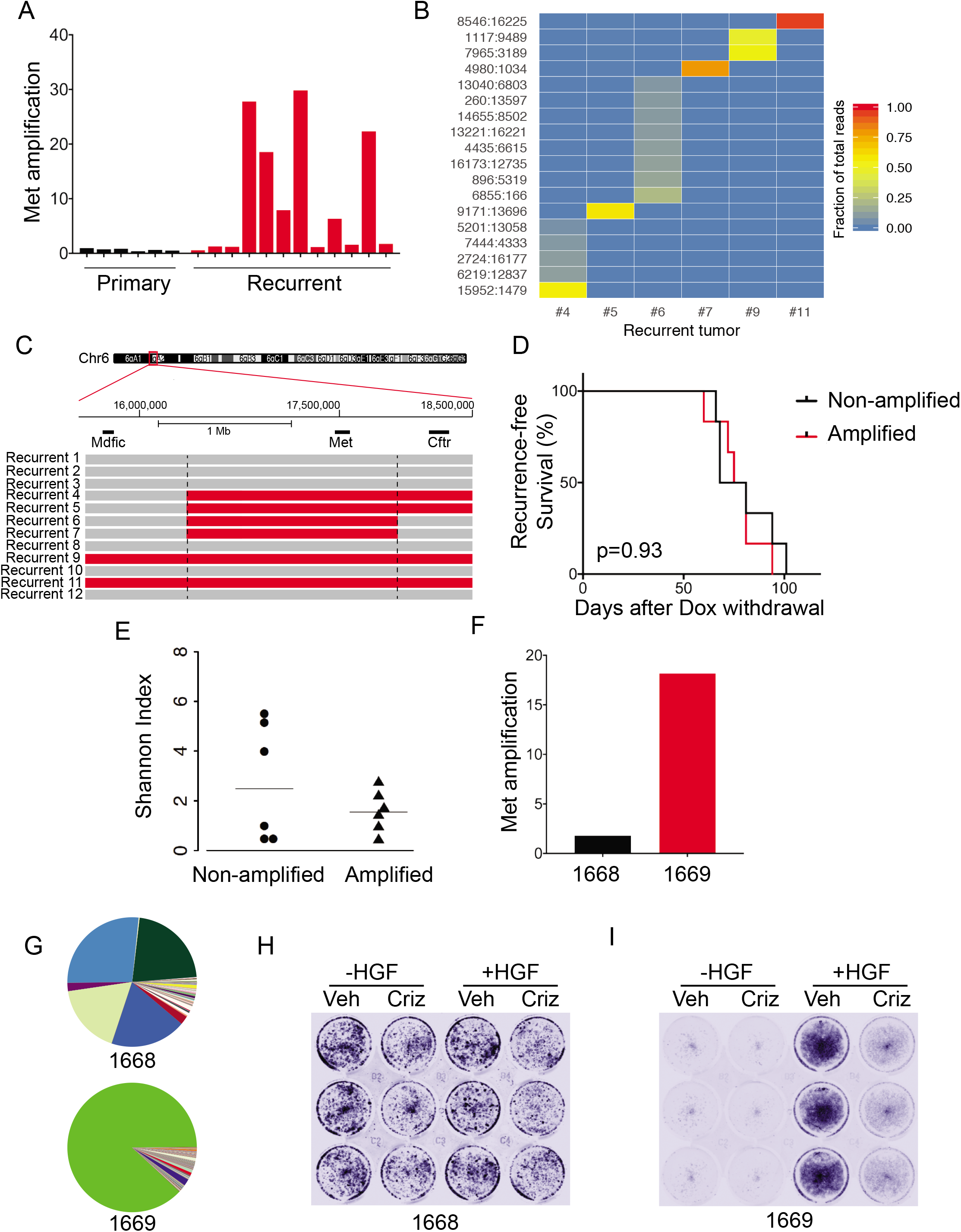
Met Amplification Drives Recurrence in a Subset of Clonal Recurrent Tumors. (A) qPCR analysis of Met copy-number in primary and recurrent tumors. Data are expressed as fold-increase in Met copy number relative to blood. (B) Heatmap of the abundance of selected barcodes in recurrent tumors, demonstrating that individual Met-amplified tumors have unique barcodes. All barcodes present at greater than 5% in any tumor are shown. (C) Met-amplified recurrent tumors have distinct Met amplicons. qPCR analysis was used to measure Mdfic, Met, and Cftr copy number in individual Met-amplified tumors. Amplified regions are shown in red. (D) Kaplan-Meier recurrence-free survival curves for orthotopic tumors, stratified based on Met amplification status. Differences in survival were calculated using the Kaplan-Meier estimator. (E) Shannon Diversity Index showing barcode complexity in recurrent tumors with and without Met amplification. (F) Met copy number in two cell lines derived from orthotopic recurrent tumors with or without Met amplification. (G) Pie charts showing the relative frequency of barcodes in cells from (F). (H-I) Met signaling drives tumor cell proliferation in Met-amplified recurrent tumor cells. Recurrent tumor cells 1668 (H) or 1669 (I) were grown in the presence of HGF and/or crizotinib for 3 days. Cells were stained using crystal violet.

We next determined whether Met amplification correlated with clonal diversity. All Met-amplified tumors had low clonal diversity (Figure 5E), consistent with the finding that these tumors all had a small number of abundant clones (see Figure 4A and 5B). Met was not amplified in any of the high-diversity tumors, or in any tumors whose clonal composition was highly correlated with primary tumors (Figure 5E and 4D). Taken together, these results suggest that distinct Met-amplified clones – possibly arising de novo – drive tumor recurrence in a subset of tumors, yielding (oligo)clonal recurrences with distinct clonal architecture from primary tumors.

We next tested whether Met-amplified tumors are sensitive to Met inhibitors. We injected the same starting population of barcoded cells (donor tumor #1) into a new cohort of recipient mice, and generated recurrent tumors as described above. We harvested and digested tumors to generate barcoded tumor cell cultures for in vitro analyses. We measured Met amplification and sequenced the barcodes to determine the clonal composition of these cultures. Cells cultured from recurrent tumor 1668 were not Met-amplified and were oligoclonal (Figure 5F-G), while cells cultured from tumor 1669 were Met-amplified and clonal (Figure 5F-G). We found that 1669 cells required HGF for their proliferation and were sensitive to the Met inhibitor crizotinib, while 1668 cells were insensitive to both HGF and crizotinib (Figure 5H-I). This suggests, consistent with previous findings, that Met-amplified tumors are sensitive to Met inhibitors (Comoglio et al., 2008; Comoglio et al., 2018; Engelman and Janne, 2008; Gherardi et al., 2012; Turke et al., 2010).

### Tumor recurrence is accompanied by an epithelial-to-mesenchymal transition

To gain insight into the mechanism underlying reactivation in polyclonal recurrent tumors, we wanted to compare gene expression profiles of primary tumors, Met-amplified recurrent tumors, and polyclonal recurrent tumors. We could not perform gene expression analysis on the original cohort of tumors, since the entire tumor sample was used to make genomic DNA. Therefore, we generated a separate cohort of primary (n=4) and recurrent barcoded (n=8) tumors using a second digested MTB;TAN tumor (donor tumor #2), and isolated both DNA and RNA from these tumors. Barcode sequencing revealed that the pattern of changes in the clonal composition of these tumors was similar to the original cohort, with a progressive decrease in the complexity of barcodes during recurrence (Supplemental Figure 4A-C). Similarly, half (n=4) of the recurrent tumors (recurrent tumors #1, 2, 3 and 6) had Met amplification (Supplemental Figure 4D), and these tumors were either clonal or oligoclonal (Supplemental Figure 4B). In contrast, tumors lacking Met amplification (recurrent tumors 4, 5, 7 and 8) were predominantly polyclonal (Supplemental Figure 4B).

Using this new cohort, we examined how gene expression patterns differed between primary tumors, Met-amplified recurrent tumors, and non-Met amplified polyclonal recurrent tumors. It has previously been shown that recurrent tumors arising in MTB;TAN mice have undergone an epithelial-to-mesenchymal transition (EMT), and experimental induction of EMT by expression of the transcription factor Snail accelerates recurrence in this model (Moody et al., 2005). Consistent with this, EMT can promote resistance to both targeted and chemotherapies (Sequist et al., 2011; Shibue and Weinberg, 2017). We therefore determined whether clonal Met-amplified recurrent tumors and/or polyclonal non-Met amplified recurrent tumors had gene expression changes suggestive of EMT. We found that all recurrent tumors had downregulated expression of the epithelial markers E-cadherin and Epcam, and upregulated the mesenchymal marker Ddr2, as compared to primary tumors (Figure 6A); these genes have been proposed as key EMT markers in vivo (Zeisberg and Neilson, 2009). Similarly, E-cadherin expression was downregulated, and the mesenchymal gene Vimentin – an in vitro marker of EMT (Zeisberg and Neilson, 2009) – was upregulated in both 1668 and 1669 recurrent tumors cells as compared to cells derived from primary donor tumors (Supplemental Figure 5A-B). These results indicate that all recurrent tumors had undergone EMT, irrespective of their clonality or Met amplification status.

**Figure 6.**
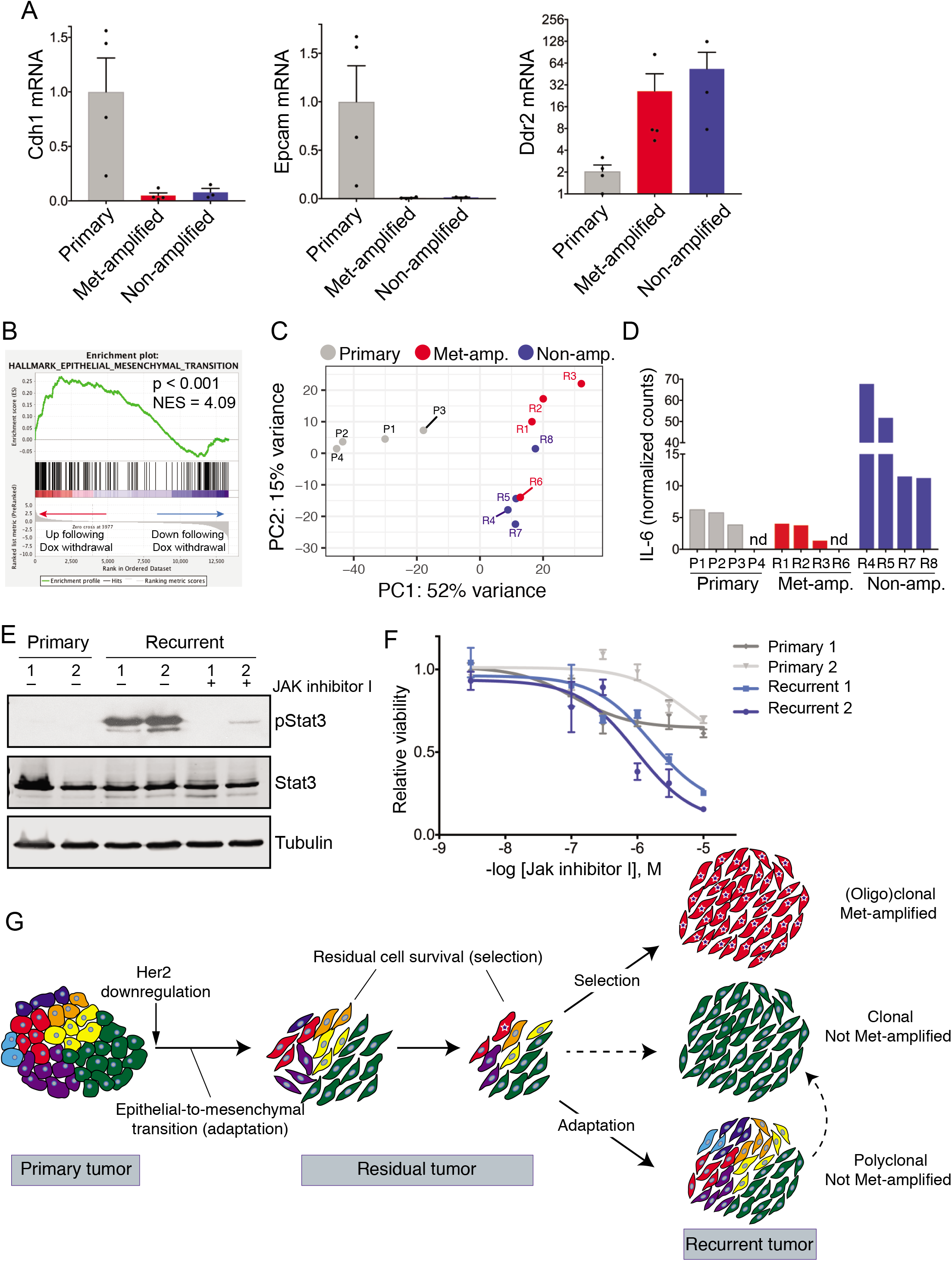
Polyclonal Recurrent Tumors Have Distinct Gene Expression Profiles. (A) qRT-PCR analysis showing expression of epithelial (Cdh1 and Epcam) and mesenchymal (Ddr2) markers in primary tumors, Met-amplified recurrent tumors, and non-amplified recurrent tumors. (B) Gene set enrichment analysis showing enrichment of an EMT signature in cells following Her2 downregulation in vitro. (C) Principal Components Analysis (PCA) of gene expression profiles in primary tumors, and recurrent tumors with and without Met amplification. (D) Normalized counts of IL-6 expression from RNA-seq analysis of primary tumors, Met-amplified recurrent tumors, and non-amplified recurrent tumors. (E) Western blot showing Stat3 phosphorylation (Y705) in primary and recurrent tumor cells derived from the autochthonous model. Note that the primary cells are donor tumor #1 and 2, and both recurrent tumor cells do not have Met amplification. (F) Dose-response curves of primary and recurrent tumor cells treated with increasing concentrations of Jak inhibitor I. (G) Model showing adaptation and selection shaping the clonal composition of tumors during residual disease and recurrence. Error bars denote mean ± SEM.

We reasoned that two mechanisms could explain this finding: recurrence could select for pre-existing mesenchymal cells in primary tumors, or EMT could be an adaptive response to Her2 inhibition. The finding that even polyclonal recurrent tumors had undergone EMT argued against a selection for pre-existing clones, since the clonal composition of polyclonal recurrent tumors resembled primary tumors. To directly assess whether EMT is an adaptive response to Her2 inhibition, we cultured cells derived from a primary tumor and removed dox to induce Her2 downregulation in vitro. We then performed RNA-seq to identify transcriptional changes 4 days following Her2 downregulation, and performed gene set enrichment analysis (GSEA) to identify gene sets enriched in cells with and without Her2 expression. The top gene set enriched in cells grown in the presence of dox was an E2F target gene set (Supplemental Figure 5C), consistent with the notion that Her2 drives proliferation of these cells (Moody et al., 2002), and thereby providing validation of this approach. We found that the top-scoring gene set enriched in the cells following dox withdrawal was an EMT signature (Figure 6B). This is consistent with the model that EMT is an adaptive response to Her2 downregulation that occurs in the absence of selection, providing a basis for the observation that all recurrent tumors have undergone EMT.

### Polyclonal recurrent tumors have distinct gene expression profiles

We next performed RNA-seq on tumors from this cohort to more broadly compare gene expression patterns between primary tumors, Met-amplified recurrent tumors, and non-Met amplified polyclonal recurrent tumors. Principal components analysis revealed that these tumors clustered by group, indicating that each of these cohorts had a distinct gene expression pattern (Figure 6C). Principal component 1 (PC1) efficiently separated primary tumors from recurrent tumors, while PC2 partly separated Met-amplified from non-amplified tumors (Figure 6D). Taken together, this suggests that polyclonal recurrent tumors have a unique gene expression profile.

The polyclonal nature of these tumors suggested that recurrence was associated with reactivation of a large number of clones in residual tumors. We reasoned that this may have been driven by a secreted factor(s) that acted in an autocrine or paracrine manner to induce tumor cell proliferation. We therefore examined the RNA-seq data for cytokines or growth factors expressed at higher levels in polyclonal recurrent tumors. We found that IL-6 was expressed between 4.9-fold and 29-fold higher in non-amplified recurrent tumors compared to Met-amplified recurrent tumors (Figure 6D). IL-6 exerts its effects by signaling through Jak kinases and Stat transcription factors (Yu et al., 2014). We therefore examined whether there was evidence for Jak-Stat pathway activation in recurrent tumors lacking Met amplification. Because we did not have protein lysates available for this cohort of tumors, we examined tumor cells cultured from Met-amplified or non-amplified autochthonous recurrent tumors arising in MTB;TAN mice. We found that levels of pStat3 were elevated in cells from nonamplified recurrent tumors, and this could be inhibited by a Jak kinase inhibitor (Figure 6E). Finally, we tested whether tumor cells from non-amplified recurrent tumors were sensitive to a Jak inhibitor. Treatment with a pan-Jak kinase inhibitor had a modest effect on primary cells, but nearly completely inhibited the growth of recurrent tumor cells (Figure 6F). These results suggest that polyclonal recurrent tumors are driven in part by autocrine or paracrine IL-6 production that signals through the Jak-Stat pathway to drive the proliferation of these cells.

## Discussion

Most deaths from breast cancer are caused by the survival and eventual recurrence of residual tumor cells after therapy (Klein, 2011). Understanding the pathways that govern residual cell survival, and how these cells resume proliferation, may suggest strategies for targeted approaches to prevent or delay relapse. As a first step in this direction, it is important to understand the clonal evolution of tumors during residual disease and recurrence; that is, to what extent does the clonal composition of residual and recurrent tumors resemble that of primary tumors? While next-generation sequencing has provided insights into clonal evolution in recurrent tumors in humans (Yates et al., 2017), the difficulty in identifying and sampling residual tumors has impeded progress in studying this critical stage. In the current study, we used DNA barcoding to directly monitor clonal evolution during tumor growth, regression, residual disease, and recurrence.

### Distinct routes to recurrence

The clonal composition of recurrent tumors suggested that recurrence can proceed through several distinct routes (Figure 6G). One subset of recurrent tumors was clonal or oligoclonal, exhibited amplification of c-Met, and was sensitive to Met inhibitors. In these tumors, recurrence is likely driven by the expansion of a Met-amplified clone. Interestingly, the finding that each Met-amplified tumor had a unique barcode and distinct amplicon boundaries suggests that these tumors arose from independent Met-amplified clones. This may argue against the presence of a single pre-existing clone with Met amplification, and instead suggest that amplification of Met occurs *de novo,* either during orthotopic tumor growth or at the residual disease stage. It is important to note, however, that we cannot rule out the possibility that the donor primary tumor had many independent Met-amplified clones, each of which was labeled with a different barcode. Notwithstanding the timing of Met amplification, these results demonstrate that Met amplification is an escape route for tumors following oncogene inhibition, consistent with findings from a number of other groups (Engelman and Janne, 2008; Engelman et al., 2007; Liu et al., 2011; Sequist et al., 2011; Turke et al., 2010).

A second subset of tumors was polyclonal; these tumors were composed of many hundreds of clones, and the distribution of these clones was similar to primary tumors. These tumors lacked Met amplification and were not sensitive to Met inhibitors. In these tumors, recurrence was likely driven by the tumor-wide reactivation of all or most of the clones present in residual tumors, resulting in recurrent tumors with a clonal architecture that resembled primary tumors. It is possible that in these tumors reactivation of residual tumor cells was mediated by a secreted factor that functions in a paracrine manner, acting on all clones. Consistent with this, polyclonal recurrences expressed high levels of IL-6 and Stat3 activation, and were sensitive to a Jak inhibitor. Interestingly, an IL-6-driven paracrine loop has been shown to drive the survival of residual lymphoma cells following chemotherapy in the thymus (Gilbert and Hemann, 2010). Our results extend this finding by suggesting that, in some cases, tumor cell-derived IL-6 may act in an autocrine manner to drive the proliferation of residual tumor cells.

In the final group of tumors, a single clone marked by barcode 8240:8518 composed nearly the entire tumor. This suggests that this clone, which was the most abundant clone present in all primary and residual tumors, expanded to give rise to recurrent tumors. These tumors were not Met-amplified, suggesting that an alternative pathway drives relapse in these tumors. It will be interesting to elucidate the event that triggers reactivation of this clone.

### Changes in clonal complexity during residual disease

Her2 downregulation and tumor regression were accompanied by a progressive decrease in the clonal diversity of tumors, consistent with the notion that only a subset of tumor cells is capable of surviving oncogene downregulation. Interestingly, we found that the residual disease stage itself is also accompanied by a decrease in the number of clones and their diversity within tumors. This suggests that, rather than being a static state, residual disease is a dynamic process with ongoing changes in clonal composition. Indeed, the reduction in diversity between early and late residual disease was as large as the reduction that accompanied tumor regression, suggesting that the magnitude of selective pressures during the residual disease stage was nearly equivalent to the selective pressure exerted by oncogene inhibition. While the cellular stresses underlying this selection remain unknown, this finding suggests that recurrent tumors may arise from a subset of residual cells that can overcome these stresses. Our findings are similar to a study that assessed clonal heterogeneity of disseminated residual tumor cells in humans (Klein et al., 2002). In this study, the authors showed that residual tumors comprise a heterogeneous mixture of clones, while clinically evident metastases seemed to result from the expansion of a subset of these clones. This suggests that the clonal changes that accompany local residual disease and recurrence in our mouse model mirror, at least in part, what is observed in disseminated residual disease and metastatic relapse in humans.

### Adaptive and selective changes during recurrence

Changes in the clonal composition of tumors at different stages of recurrence suggested that both adaptive and selective pressures act to drive tumor evolution and relapse. All tumors – irrespective of their clonality or Met amplification status – had gene expression patterns consistent with EMT. This suggests that EMT did not result from the selection of a subset of clones with mesenchymal characteristics, since the clonal composition of polyclonal recurrent tumors was very similar to primary tumors. Instead, these results indicate that EMT may be an adaptive response to oncogene inhibition. Gene expression changes in tumor cells shortly following Her2 downregulation are consistent with this. It is important to note, however, that experimental induction of EMT accelerates recurrence in this model (Moody et al., 2005). Taken together, these findings suggest that EMT is an adaptive response that promotes cell survival following oncogene inhibition, but that the ability to undergo a full EMT is nonetheless rate-limiting for recurrence, perhaps because mesenchymal cells can better survive oncogene downregulation. Consistent with this, residual breast cancer cells that can survive neoadjuvant therapy display mesenchymal characteristics (Creighton et al., 2009). More broadly, the observation that some recurrent tumors had very similar clonal composition to primary tumors is *prima facie* evidence that recurrence in this subset of tumors was driven by adaptive mechanisms. In contrast, Met-amplified recurrent tumors usually comprised one or two clones, suggesting that recurrence in these tumors proceeds via selection. Together, these results indicate that adaptation and selection can work together to shape the evolution of tumors during residual disease and recurrence.

The findings presented here are reminiscent of results describing the emergence of resistance to EGFR inhibitors in EGFR-mutant lung cancer (Hata et al., 2016). That study found that resistance could emerge either from the expansion of cells with pre-existing resistance mutations, or through the survival of a population of drug-tolerant persister cells. In this latter case, drug-tolerant cells can survive but not proliferate in the presence of EGFR inhibitors; these cells eventually evolve mutations rendering them fully resistant. In our study, we find that residual cells are capable of surviving Her2 downregulation but are not initially competent to proliferate, suggesting that these cells may be analogous to persister cells. A unique finding in our study is that, in some instances, the majority of these residual cells can acquire the ability to resume proliferation in the absence of additional clonal selection. This represents a novel mode of recurrence in which some signal – perhaps a secreted paracrine-acting factor such as IL-6 – can induce the reactivation of residual tumor cells *en masse.* Approaches to target this resistance mechanisms may block this route of recurrence.

## Methods

### Cell culture, barcode library transduction, and dose-response curves

Primary tumor cells from two independent MTB;TAN primary tumors were used for all experiments and have been described previously (Mabe et al., 2018). These cells were cultured in the presence of dox to maintain Her2 expression and used at early passage number (prior to passage 15). To generate barcoded populations, cells were infected with the CellTracker lentiviral barcode library from Cellecta, which contains approximately 50 million unique barcodes. 200,000 cells were transduced with this library in the presence of polybrene (8 μg/ml) at an MOI of 0.1 to yield a population with approximately 20,000 unique barcodes. Cells were expanded for 12 population doublings prior to injection into mice.

Cell lines were generated from barcoded recurrent tumors using a previously described protocol (Mabe et al., 2018). Briefly, tumors were harvested, minced, and digested in collagenase and hyaluronidase (StemCell Technologies). Digestion media was removed and the cells were resuspended in red blood cell lysis buffer, followed by Dispase II (5 mg/mL) and DNase I (100 μg/ml) prior to plating. Cells were grown in DMEM with 10% SCS, 1% Pen-Step, 1% Glutamine, and supplemented with EGF (0.01 μg/ml, Sigma) and insulin (5 μg/ml, Gemini Bioproducts). To generate Met-amplified cell cultures, cells were supplemented with HGF (250 ng/ml, R&D systems). Crizonitib was used at 500 nM (Selleck). Recurrent tumor cells derived from recurrent tumors arising in the autochthonous MTB;TAN mouse model were cultured as previously described (Mabe et al., 2018).

Dose response curves were performed as previously described (Mabe et al., 2018). Briefly, 1000 primary or recurrent tumor cells were plated onto 96-well plates. The following day, media containing 1% serum with increasing concentrations of Jak Inhibitor I (Calbiochem) was added to cells. Cell viability was measured 3 days later using CellTiterGlo (Promega).

For RNA-seq analysis, primary Her2-driven tumor cells were cultured as mammospheres (Dontu et al., 2003) in the presence of 2 μg/ml doxycycline. To induce Her2 downregulation, dox was removed from cultures, and RNA was harvested 4 days later.

### Mice

All experiments were approved by Duke IACUC (Approval #A199-17-08). One million barcoded MTB;TAN tumor cells were injected into the inguinal mammary gland of nu/nu mice. Mice were provided dox (2 mg/ml, 5% sucrose) in their drinking water two days prior to injection and remained on dox during the course of primary tumor growth. Mice were palpated twice per week to monitor growth. Once tumors reached 5 mm in diameter, one cohort of mice was sacrificed, and dox was removed from the drinking water of the other cohorts to initiate Her2 downregulation. Additional cohorts were sacrificed at 4 weeks and 8 weeks following dox withdrawal. A final cohort was monitored for the appearance of recurrent tumors, and mice were sacrificed when tumors reached between 8 and 12 mm in diameter. Primary and recurrent tumors were harvested from mice and snap frozen for DNA isolation. Residual tumors were microdissected with the aid of a fluorescent microscope and snap frozen.

### DNA isolation, library preparation and barcode sequencing

To isolate genomic DNA, tumors were digested with Proteinase K in Buffer ATL (Qiagen). Following digestion, cells were lysed by adding SDS (final concentration, 0.5%) and sonicating to shear DNA. Genomic DNA was then purified using phenol-chloroform extraction followed by isopropanol precipitation.

To prepare libraries for sequencing, DNA concentration was measured using Broad Range Qubit methodology (Invitrogen). 80 ng of DNA (approximately 12,100 genomes) was used as input for each single reaction. To increase the number of analyzed cells we performed 8 reactions per sample; therefore, barcodes from around 97,000 cells were analyzed for each sample. We used a two-step PCR amplification protocol as described in Lundberg *et. al.* (Lundberg et al., 2013). In the first step of the reaction barcodes were amplified using custom primers (Forward: “Step1_F”, Reverse: “Step1_R”) for 10 cycles. KAPA Robust 2G polymerase (KAPA Biosystems-Roche) was used during this step to ensure efficiency and specificity of barcode amplification. DNA was purified between steps one and two using magnetic beads at a 1.3(beads):1(reaction) proportion. During the second step, PCR products from step one were amplified using custom primers (Forward: “ClonalBarcode Adaptor_1”, Reverse: “PCR Primer_IndexN”) for 23 cycles. KAPA Hifi Polymerase (KAPA Biosystems-Roche) was used during this step to ensure fidelity of the amplification. DNA was purified after the second step using magnetic beads at a 1(beads): 1(reaction) proportion. Amplified barcodes were sequenced using custom “Clonal Barcodes R1” sequencing primer and standard TruSeq sequencing primers using an Illumina HiSeq2500 Rapid Single End 40×9. Custom primer sequences are shown in Supplemental Table 1 and 2.

### RNA-seq, qRT-PCR, copy-number analysis, and western blotting

RNA was isolated from cells or tumors using the RNeasy kit (Qiagen). RNA was sequenced using the Illumina HiSeq 4000 libraries and sequencing platform with 50 base pair single end reads by the Duke GCB Sequencing and Genomic Technologies Shared Resource (Durham, NC).

qRT-PCR was performed as described (Mabe et al., 2018) using the following TaqMan probes: Cdh1 (Mm01247357_m1), Epcam (Mm00493214_m1), Ddr2 (Mm00445615_m1), and Vim (Mm01333430_m1) and normalized to expression of Actb (Mm02619580_g1).

To measure the copy number of Met and flanking genes, we performed real-time PCR on purified genomic DNA using the following TaqMan probes (all from Applied Biosystems): Met (Mm00565151_cn), Mdfic (Mm00565127_cn), Cav2 (Mm00181583_cn), and Cftr (Mm00181608_cn). Absolute copy-number for each gene was calculated by normalizing to an internal reference gene (Tfrc) and expressed as fold-change over genomic DNA from non-tumor tissue.

Western blotting was performed as previously described (Damrauer et al., 2018) using antibodies against total Stat3 (Cell Signaling), phospho-Stat3 Y705 (Cell Signaling), and tubulin (Sigma).

### Bioinformatics

#### Barcode mapping

Low quality reads with mean Phred scores smaller than 30 were excluded from the analysis. Remaining reads with a length of 40 base pair (bp) were split into left and right short reads (18 bp) after removing the 4 bp linkers at the center. The resulted short left and right RNA reads were mapped to the sequencing library. The mapping allowed up to 2 mismatches. After mapping, the mapped results for short left and right RNA reads were combined. Based on the combined mapping results, a count table listing the number of reads for each barcode in each sample was generated for the further statistical analysis. Barcodes with read counts smaller than 1 were excluded from downstream statistical analysis. Numbers of unique barcodes, proportions of reads for the barcodes, most abundant sequenced barcodes, and Shannon diversity indexes for each sample were obtained based on the barcode reads count table. Dissimilarities based on Jensen-Shannon divergence were obtained to evaluate the differences in clonal composition among different tumors.

#### RNA-Seq analysis

FastQC (v0.11.5) (Andrews, S., FastQC A Quality Control tool for High Throughput Sequence Data. 2014.; https://www.bioinformatics.babraham.ac.uk/projects/fastqc/) was used to assess the general sequencing quality. All the samples were sequenced at a length of 51bp. All the samples have an averaged Phred score greater than 30. RNA sequences which passed quality control were aligned to reference mouse genome GRCm38 (mm10) using STAR (v2.5.2b) (Dobin et al., 2013). The aligned reads were mapped to genomic features using HTSeq (Anders et al., 2015), implemented in the STAR program. The reference sequence and GTF file were obtained from the NCBI GRCm38 bundle available from the iGenomes collection. Gene differential expression was performed within the framework of a negative binomial model using R (v3.4.4) (R Core Team, R: A Language and Environment for Statistical Computing. 2016: Vienna, Austria.) and its extension package DESeq2 (v1.18.1) (Anders and Huber, 2010). PCA analysis were performed for each of the cohort based on the normalized reads matrix (based on rlog from DESeq2(v1.18.1)) from the DESeq2 (v1.18.1).

### Statistics

A t-test was used to test the changes of the number of unique barcodes, the Shannon diversity index, the number of the most abundant barcodes that composing 50% of the total reads, and the most abundant barcode proportion for injected cells, primary tumors, and residual tumors. t-test P-values were reported. Two-way ANOVA test was used to test the mean changes of Shannon diversity indexes from primary tumors to 4 week residual tumors and 4 week residual tumors to 8 week residual tumors. Unadjusted P-value of the interaction term from two-way ANOVA test was reported. Cox proportional hazard regression was used for the time to tumor recurrence analyses (Cox and Oakes, 1984) using the R extension package survival v2.41-3 (Therneau T., A Package for Survival Analysis in S. version 2.38, 2015; https://CRAN.R-project.org/package=survival). The Cox score statistics (Cox and Oakes, 1984) was used to test the difference of time to tumor recurrence between Met amplified and non-Met amplified tumors. The P-values were not adjusted for multiple testing. The Kaplan-Meier product estimator was used to estimate the survival function for the tumor recurrence for both Met amplified and non-Met amplified recurrent tumors.

### Programming and documentation

Statistical analyses were mainly scripted using the R statistical environment[R] along with its extension packages from the comprehensive R archive network (CRAN; https://cran.r-project.org/) and the Biocoductor project[BIOC] (Gentleman et al., 2004). All analyses were programmed and documented using mercurial (https://www.mercurial-scm.org/) for source code management in Bitbucket (https://bitbucket.org/product).

## Data availability

All sequencing data, including barcode sequencing and RNA sequencing, are available at Sequence Read Archive (https://www.ncbi.nlm.nih.gov/Traces/study/?acc=PRJNA509416). Code for barcode mapping and data analysis is available at https://bitbucket.org/dcibioinformatics/alvarez-paper-submit.

## Acknowledgements

We thank Nicolas Devos from the Duke Sequencing and Genomic Technologies Core for RNA-seq library preparation and sequencing. We thank Daniel Abravanel and David MacAlpine for helpful discussions and critical reading of the manuscript. This work was funded by the National Cancer Institute under award numbers R01CA208042 (to JVA) and F31CA220957 (to AW) and by startup funds from the Duke Cancer Institute, the Duke University School of Medicine and the Whitehead Foundation (to JVA).

## Author Contributions

J.V.A., A.W., and J.S.D. were responsible for the conception, design, and interpretation of all experiments. A.W., J.S.D., R.L., R.N., D.B.F., and N.M. performed experiments and collected data. T.D., H.K., and P.M. performed the barcode library preparation and sequencing. J.L. and J.G designed and performed the bioinformatics and statistical analysis, and K.O. supervised all bioinformatics and statistical work. A.W., J.L., K.O. and J.V.A. wrote the manuscript. J.V.A. supervised all work.

## Supplemental Figure Legends

<u>Supplemental Figure 1</u>

(A) Barcode abundance in primary tumors.

(B) Barcode abundance in early residual tumors (4 weeks following dox withdrawal).

(C) Barcode abundance in late residual tumors (8 weeks following dox withdrawal).

For all plots, barcodes are ranked on the x-axis from most to least abundant. Red line denotes 10% abundance.

<u>Supplemental Figure 2</u>

The frequency of Barcode 8240:8518 in primary, residual, and recurrent tumors. Individual recurrent tumors are labeled, and Met-amplified recurrent tumors are shown in red.

<u>Supplemental Figure 3</u>

(A) – (G) The number of reads across all tumors for the most abundant barcodes in recurrent tumors #4-9 and 11.

<u>Supplemental Figure 4</u>

(A) An independent donor tumor (donor tumor #2) was infected with the barcode library and injected into mice as in Figure 1. Pie charts show barcode abundance in the injected cell population and 4 independent primary tumors.

(B) Pie charts showing the relative frequency of barcodes in 8 independent recurrent tumors.

(C) Shannon diversity index showing barcode complexity of the starting cell population, 4 primary tumors, and 8 recurrent tumors.

(D) qPCR analysis of Met copy number in primary and recurrent tumors. Data are expressed as fold-increase in Met copy number relative to blood.

<u>Supplemental Figure 5</u>

(A) – (B) qRT-PCR analysis showing expression of epithelial (Cdh1) and mesenchymal (Vim) markers in cells derived from primary tumors (donor tumor #1 and 2), or from orthotopic recurrent tumors with (1669) or without (1668) Met amplification.

(C) Gene set enrichment analysis showing enrichment of an E2F signature in cells grown in the presence of Dox (Her2 on).

## References

Abravanel, D. L., Belka, G. K., Pan, T. C., Pant, D. K., Collins, M. A., Sterner, C. J., and Chodosh, L. A. (2015). Notch promotes recurrence of dormant tumor cells following HER2/neu-targeted therapy. J Clin Invest 125, 2484–2496.

Acharyya, S., Oskarsson, T., Vanharanta, S., Malladi, S., Kim, J., Morris, P. G., Manova-Todorova, K., Leversha, M., Hogg, N., Seshan, V. E., et al. (2012). A CXCL1 paracrine network links cancer chemoresistance and metastasis. Cell 150, 165–178.

Alvarez, J. V., Pan, T. C., Ruth, J., Feng, Y., Zhou, A., Pant, D., Grimley, J. S., Wandless, T. J., Demichele, A., Investigators, I. S. T., and Chodosh, L. A. (2013). Par-4 downregulation promotes breast cancer recurrence by preventing multinucleation following targeted therapy. Cancer Cell 24, 30–44.

Anders, S., and Huber, W. (2010). Differential expression analysis for sequence count data. Genome Biol 11, R106.

Anders, S., Pyl, P. T., and Huber, W. (2015). HTSeq--a Python framework to work with high-throughput sequencing data. Bioinformatics 31, 166–169.

Bhang, H. E., Ruddy, D. A., Krishnamurthy Radhakrishna, V., Caushi, J. X., Zhao, R., Hims, M. M., Singh, A. P., Kao, I., Rakiec, D., Shaw, P., et al. (2015). Studying clonal dynamics in response to cancer therapy using high-complexity barcoding. Nat Med 21, 440–448.

Bivona, T. G., and Doebele, R. C. (2016). A framework for understanding and targeting residual disease in oncogene-driven solid cancers. Nat Med 22, 472–478.

Comoglio, P. M., Giordano, S., and Trusolino, L. (2008). Drug development of MET inhibitors: targeting oncogene addiction and expedience. Nat Rev Drug Discov 7, 504–516.

Comoglio, P. M., Trusolino, L., and Boccaccio, C. (2018). Known and novel roles of the MET oncogene in cancer: a coherent approach to targeted therapy. Nat Rev Cancer 18, 341–358.

Cox, D. R., and Oakes, D. (1984). Analysis of survival data, (London; New York: Chapman and Hall).

Creighton, C. J., Li, X., Landis, M., Dixon, J. M., Neumeister, V. M., Sjolund, A., Rimm, D. L., Wong, H., Rodriguez, A., Herschkowitz, J. I., et al. (2009). Residual breast cancers after conventional therapy display mesenchymal as well as tumor-initiating features. Proc Natl Acad Sci U S A 106, 13820–13825.

Damrauer, J. S., Phelps, S. N., Amuchastegui, K., Lupo, R., Mabe, N. W., Walens, A., Kroger, B. R., and Alvarez, J. V. (2018). Foxo-dependent Par-4 Upregulation Prevents Long-term Survival of Residual Cells Following PI3K-Akt Inhibition. Mol Cancer Res.

Dobin, A., Davis, C. A., Schlesinger, F., Drenkow, J., Zaleski, C., Jha, S., Batut, P., Chaisson, M., and Gingeras, T. R. (2013). STAR: ultrafast universal RNA-seq aligner. Bioinformatics 29, 15–21.

Dontu, G., Abdallah, W. M., Foley, J. M., Jackson, K. W., Clarke, M. F., Kawamura, M. J., and Wicha, M. S. (2003). In vitro propagation and transcriptional profiling of human mammary stem/progenitor cells. Genes Dev 17, 1253–1270.

Engelman, J. A., and Janne, P. A. (2008). Mechanisms of acquired resistance to epidermal growth factor receptor tyrosine kinase inhibitors in non-small cell lung cancer. Clin Cancer Res 14, 2895–2899.

Engelman, J. A., Zejnullahu, K., Mitsudomi, T., Song, Y., Hyland, C., Park, J. O., Lindeman, N., Gale, C. M., Zhao, X., Christensen, J., et al. (2007). MET amplification leads to gefitinib resistance in lung cancer by activating ERBB3 signaling. Science 316, 1039–1043.

Feng, Y., Pan, T. C., Pant, D. K., Chakrabarti, K. R., Alvarez, J. V., Ruth, J. R., and Chodosh, L. A. (2014). SPSB1 promotes breast cancer recurrence by potentiating c-MET signaling. Cancer Discov 4, 790–803.

Fuglede, B., and Topsoe, F. (2004). Jensen-Shannon divergence and Hilbert space embedding. International Symposium onInformation Theory, 2004 ISIT 2004 Proceedings, 31–.

Gentleman, R. C., Carey, V. J., Bates, D. M., Bolstad, B., Dettling, M., Dudoit, S., Ellis, B., Gautier, L., Ge, Y., Gentry, J., et al. (2004). Bioconductor: open software development for computational biology and bioinformatics. Genome Biol 5, R80.

Ghajar, C. M. (2015). Metastasis prevention by targeting the dormant niche. Nat Rev Cancer 15, 238–247.

Gherardi, E., Birchmeier, W., Birchmeier, C., and Vande Woude, G. (2012). Targeting MET in cancer: rationale and progress. Nat Rev Cancer 12, 89–103.

Gilbert, L. A., and Hemann, M. T. (2010). DNA damage-mediated induction of a chemoresistant niche. Cell 143, 355–366.

Gonzalez-Angulo, A. M., Morales-Vasquez, F., and Hortobagyi, G. N. (2007). Overview of resistance to systemic therapy in patients with breast cancer. Adv Exp Med Biol 608, 1–22.

Goss, P. E., and Chambers, A. F. (2010). Does tumour dormancy offer a therapeutic target? Nat Rev Cancer 10, 871–877.

Hata, A. N., Niederst, M. J., Archibald, H. L., Gomez-Caraballo, M., Siddiqui, F. M., Mulvey, H. E., Maruvka, Y. E., Ji, F., Bhang, H. E., Krishnamurthy Radhakrishna, V., et al. (2016). Tumor cells can follow distinct evolutionary paths to become resistant to epidermal growth factor receptor inhibition. Nat Med 22, 262–269.

Jones, S. E. (2008). Metastatic breast cancer: the treatment challenge. Clin Breast Cancer 8, 224–233.

Klein, C. A. (2011). Framework models of tumor dormancy from patient-derived observations. Curr Opin Genet Dev 21, 42–49.

Klein, C. A., Blankenstein, T. J., Schmidt-Kittler, O., Petronio, M., Polzer, B., Stoecklein, N. H., and Riethmuller, G. (2002). Genetic heterogeneity of single disseminated tumour cells in minimal residual cancer. Lancet 360, 683–689.

Liu, P., Cheng, H., Santiago, S., Raeder, M., Zhang, F., Isabella, A., Yang, J., Semaan, D. J., Chen, C., Fox, E. A., et al. (2011). Oncogenic PIK3CA-driven mammary tumors frequently recur via PI3K pathway-dependent and PI3K pathway-independent mechanisms. Nat Med 17, 1116–1120.

Lundberg, D. S., Yourstone, S., Mieczkowski, P., Jones, C. D., and Dangl, J. L. (2013). Practical innovations for high-throughput amplicon sequencing. Nat Methods 10, 999–1002.

Mabe, N. W., Fox, D. B., Lupo, R., Decker, A. E., Phelps, S. N., Thompson, J. W., and Alvarez, J. V. (2018). Epigenetic silencing of tumor suppressor Par-4 promotes chemoresistance in recurrent breast cancer. J Clin Invest 128, 4413–4428.

Magurran, A. E. (2005). Biological diversity. Curr Biol 15, R116–118.

Marusyk, A., Almendro, V., and Polyak, K. (2012). Intra-tumour heterogeneity: a looking glass for cancer? Nat Rev Cancer 12, 323–334.

Moody, S. E., Perez, D., Pan, T. C., Sarkisian, C. J., Portocarrero, C. P., Sterner, C. J., Notorfrancesco, K. L., Cardiff, R. D., and Chodosh, L. A. (2005). The transcriptional repressor Snail promotes mammary tumor recurrence. Cancer Cell 8, 197–209.

Moody, S. E., Sarkisian, C. J., Hahn, K. T., Gunther, E. J., Pickup, S., Dugan, K. D., Innocent, N., Cardiff, R. D., Schnall, M. D., and Chodosh, L. A. (2002). Conditional activation of Neu in the mammary epithelium of transgenic mice results in reversible pulmonary metastasis. Cancer Cell 2, 451–461.

Nguyen, L. V., Cox, C. L., Eirew, P., Knapp, D. J., Pellacani, D., Kannan, N., Carles, A., Moksa, M., Balani, S., Shah, S., et al. (2014). DNA barcoding reveals diverse growth kinetics of human breast tumour subclones in serially passaged xenografts. Nat Commun 5, 5871.

Nguyen, L. V., Pellacani, D., Lefort, S., Kannan, N., Osako, T., Makarem, M., Cox, C. L., Kennedy, W., Beer, P., Carles, A., et al. (2015). Barcoding reveals complex clonal dynamics of de novo transformed human mammary cells. Nature 528, 267–271.

Saphner, T., Tormey, D. C., and Gray, R. (1996). Annual hazard rates of recurrence for breast cancer after primary therapy. J Clin Oncol 14, 2738–2746.

Sequist, L. V., Waltman, B. A., Dias-Santagata, D., Digumarthy, S., Turke, A. B., Fidias, P., Bergethon, K., Shaw, A. T., Gettinger, S., Cosper, A. K., et al. (2011). Genotypic and histological evolution of lung cancers acquiring resistance to EGFR inhibitors. Sci Transl Med 3, 75ra26.

Shibue, T., and Weinberg, R. A. (2017). EMT, CSCs, and drug resistance: the mechanistic link and clinical implications. Nat Rev Clin Oncol 14, 611–629.

Sosa, M. S., Bragado, P., and Aguirre-Ghiso, J. A. (2014). Mechanisms of disseminated cancer cell dormancy: an awakening field. Nat Rev Cancer 14, 611–622.

Turke, A. B., Zejnullahu, K., Wu, Y. L., Song, Y., Dias-Santagata, D., Lifshits, E., Toschi, L., Rogers, A., Mok, T., Sequist, L., et al. (2010). Preexistence and clonal selection of MET amplification in EGFR mutant NSCLC. Cancer Cell 17, 77–88.

Yates, L. R., Knappskog, S., Wedge, D., Farmery, J. H. R., Gonzalez, S., Martincorena, I., Alexandrov, L. B., Van Loo, P., Haugland, H. K., Lilleng, P. K., et al. (2017). Genomic Evolution of Breast Cancer Metastasis and Relapse. Cancer Cell 32, 169–184 e167.

Yu, H., Lee, H., Herrmann, A., Buettner, R., and Jove, R. (2014). Revisiting STAT3 signalling in cancer: new and unexpected biological functions. Nat Rev Cancer 14, 736–746.

Zeisberg, M., and Neilson, E. G. (2009). Biomarkers for epithelial-mesenchymal transitions. J Clin Invest 119, 1429–1437.

