## Supplemental Figures for "Adaptation and Selection Shape Clonal Evolution During Residual Disease and Recurrence"

**Supplemental Figure 1**

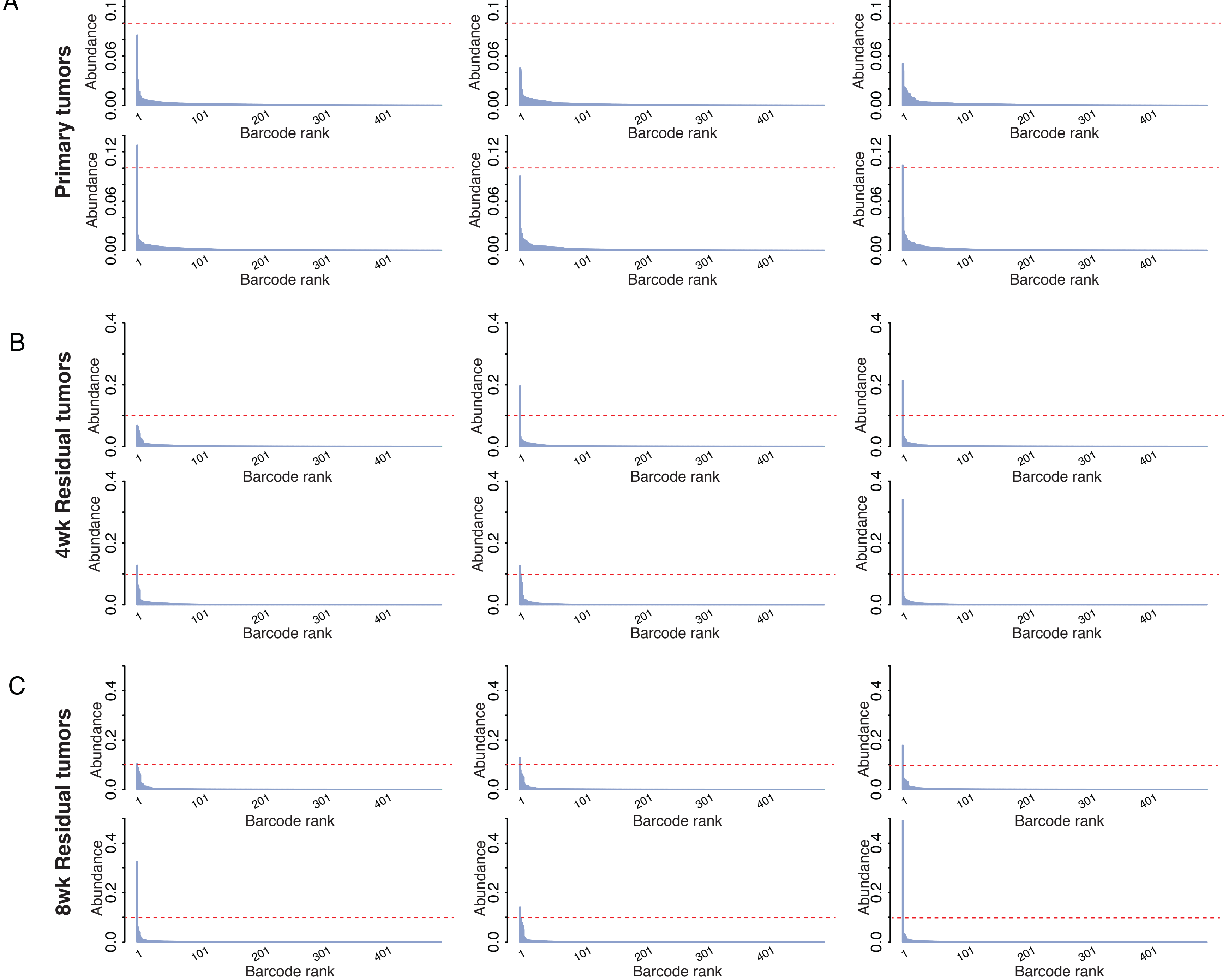

**Supplemental Figure 2**

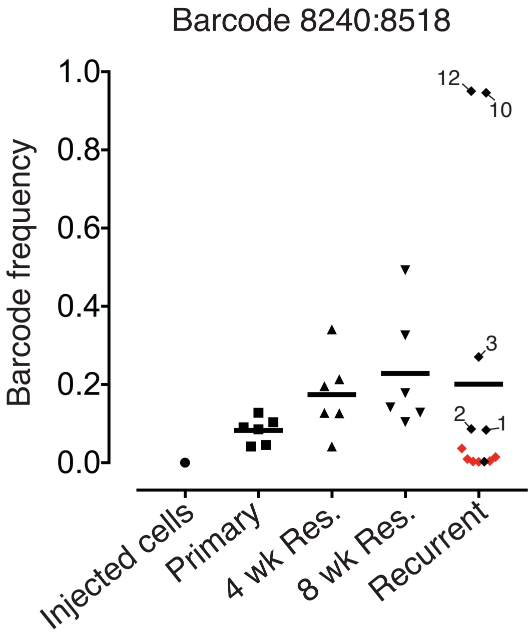

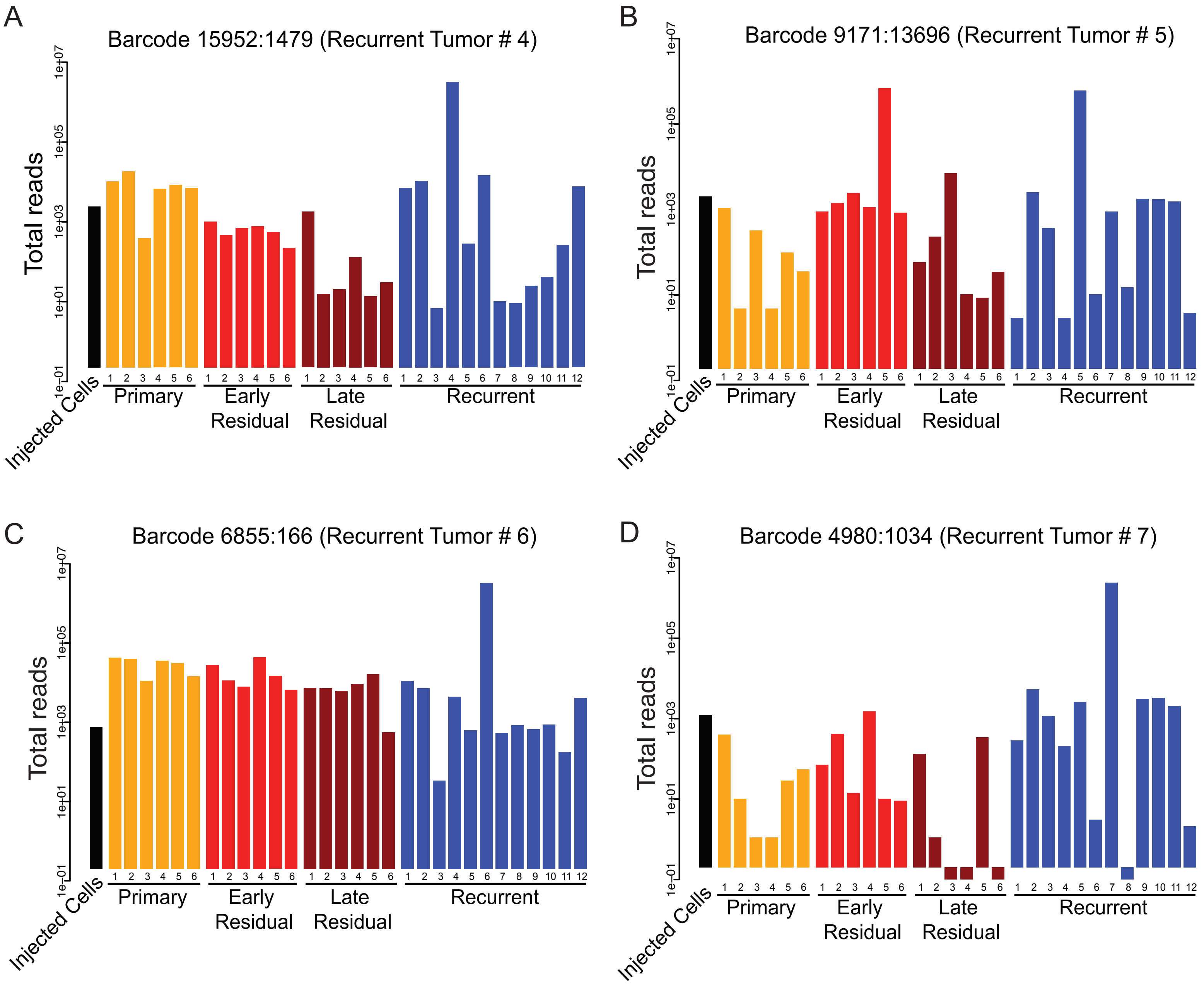
A
Barcode 15952:1479 (Recurrent Tumor # 4)
B
Barcode 9171:13696 (Recurrent Tumor # 5)
C
Barcode 6855:166 (Recurrent Tumor # 6)
D
Barcode 4980:1034 (Recurrent Tumor # 7)
Total reads
1e+07
1e+05
1e+03
1e+01
1e-01
Injected Cells
Primary
Early Residual
Late Residual
Recurrent

### Supplemental Figure 3

E

Barcode 11038:14648 (Recurrent Tumor #8)

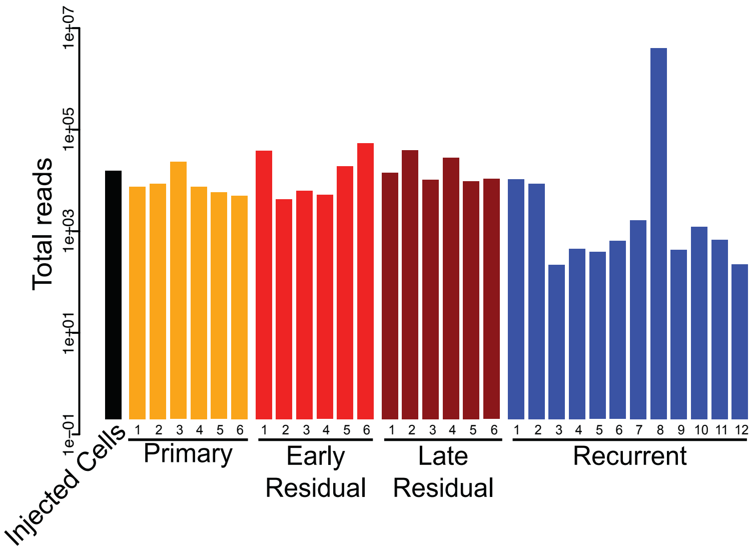


F

Barcode 7965:3189 (Recurrent Tumor #9)

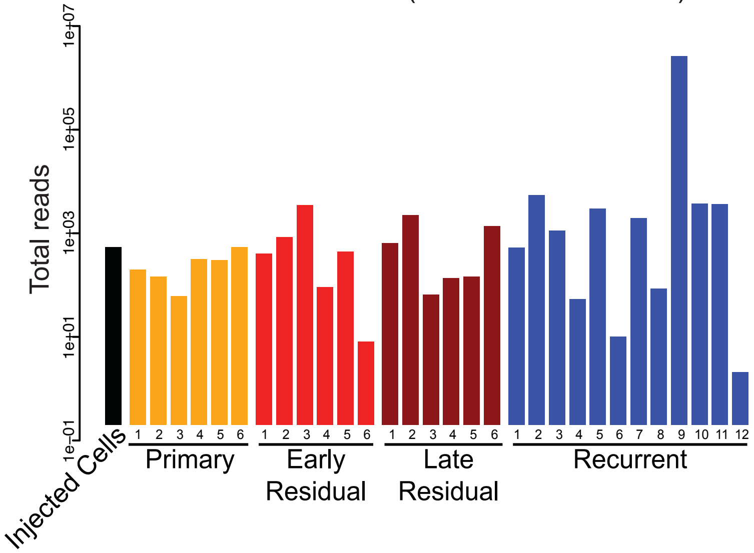


G

Barcode 8546:16225 (Recurrent Tumor #11)

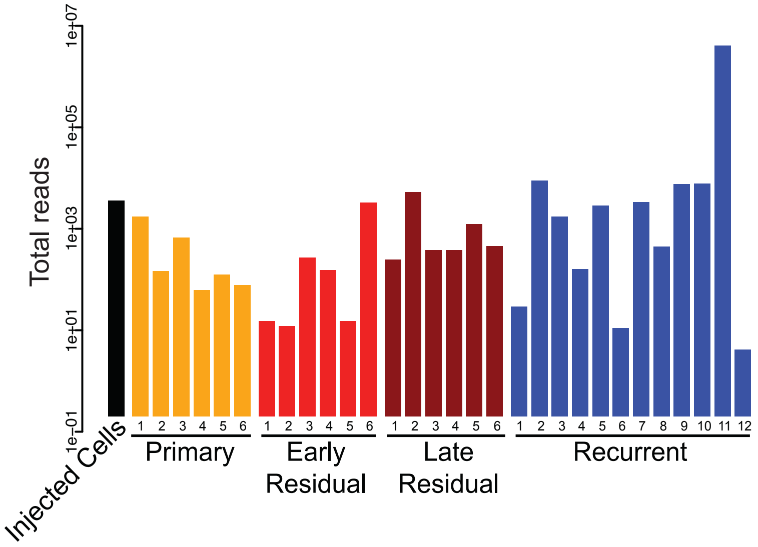

### Supplemental Figure 4

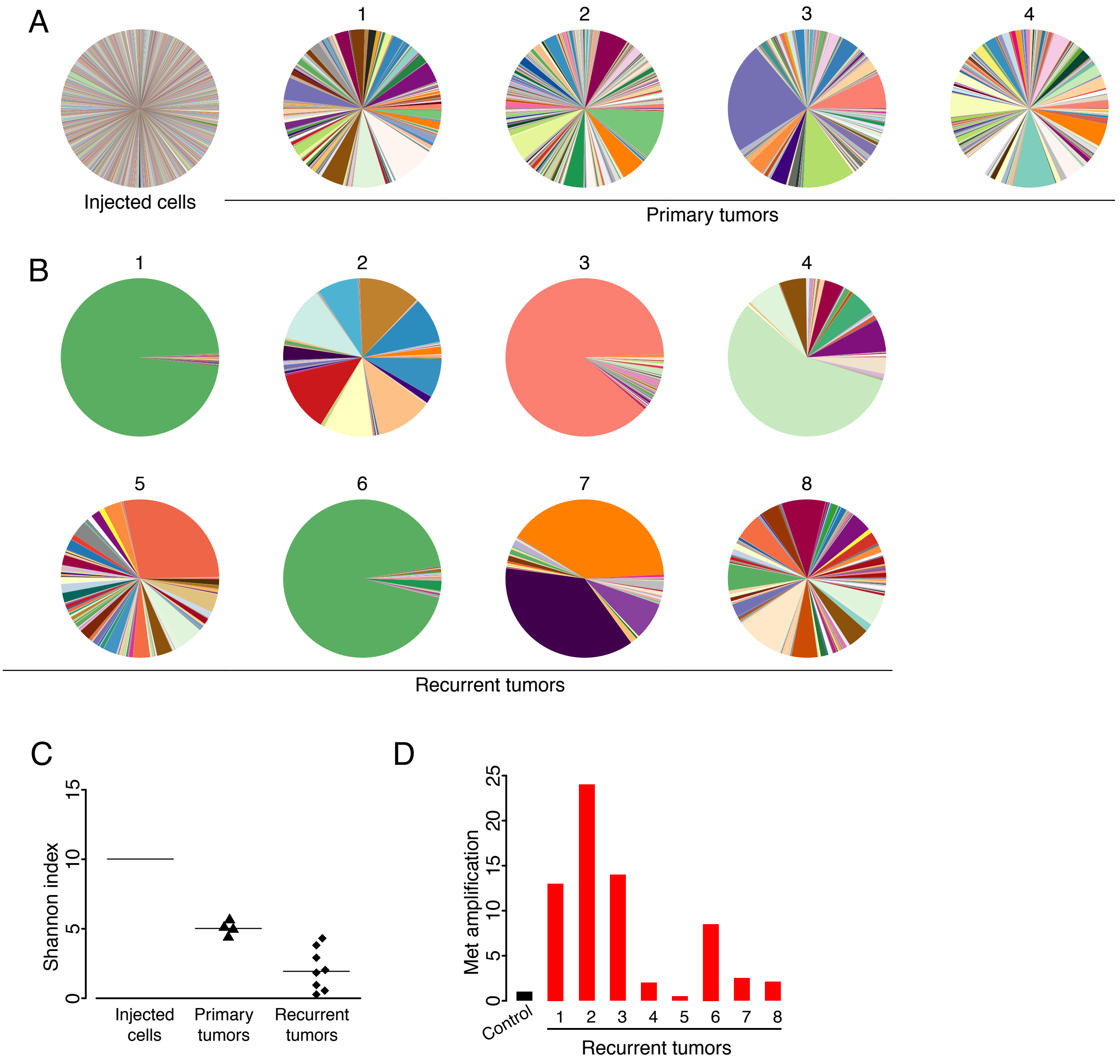

#### Supplemental Figure 5

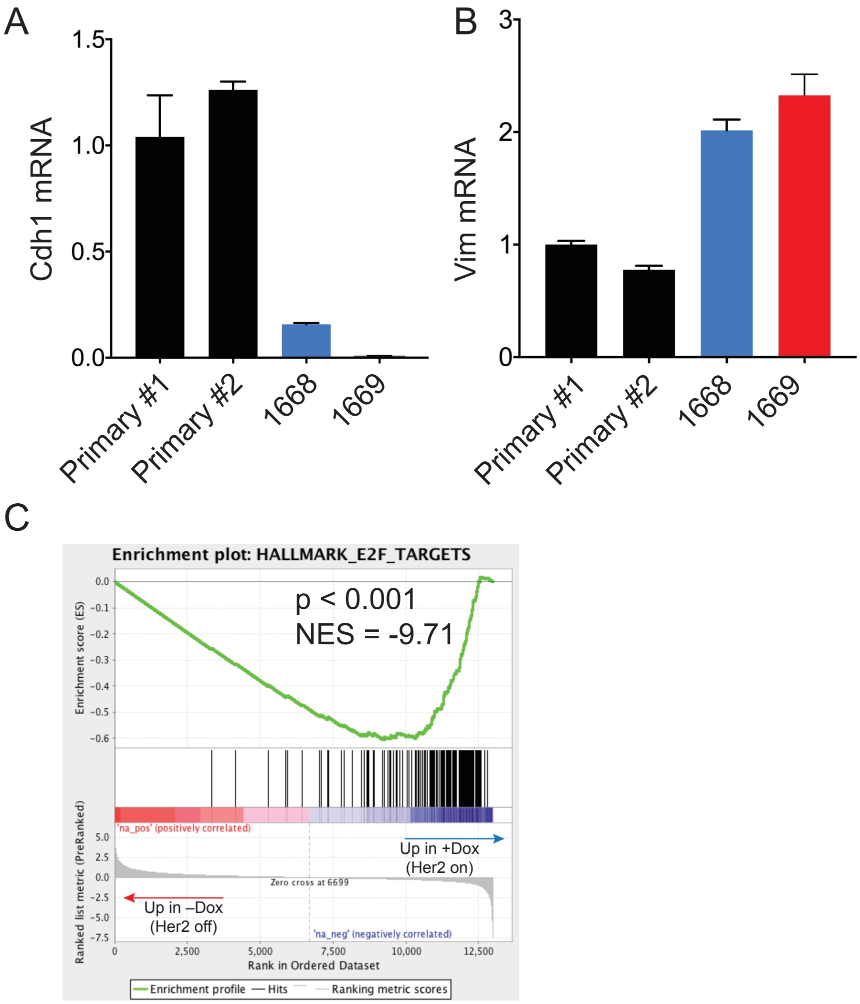
